# Dysregulation of synaptic-related genes of neuroimmune networks within peripheral blood mononuclear cells in major depressive disorder

**DOI:** 10.1101/2024.11.13.623441

**Authors:** Anny Silva Adri, Adriel Leal Nóbile, Débora Gomes de Alburquerque, Pedro Marçal Barcelos, Fernando Yuri Nery do Vale, Ian Antunes F. Bahia, Paula P. Freire, Roseane Galdioli Nava, Yohan Lucas G. Correa, Gustavo Cabral-Miranda, Rodrigo Dalmolin, Lena F. Schimke, Dennyson Leandro M. Fonseca, Igor Salerno Filgueiras, Helder I. Nakaya, Rafael Machado Rezende, Haroldo Dutra Dias, Otavio Cabral-Marques

## Abstract

Major depressive disorder (MDD) involves complex neuroimmune interactions linked to gene modulation. Our study investigates synaptic-related gene dysregulation in peripheral blood mononuclear cells (PBMCs) from MDD patients, showing how these immune cells mirror neural processes. Using RNA-seq data, we identified 1,383 differentially expressed genes (DEGs) related to neuroimmune crosstalk, with 49 DEGs effectively distinguishing MDD patients from controls based on synaptic functions. Synaptic genes, enriched for roles like vesicle transport, suggest mechanistic links between immune cells and neural signaling. Eleven synaptic-related DEGs were shared between PBMCs and brain regions involved in mood regulation, highlighting a common molecular signature. Among them, *ADORA3* and *RPS28* emerged as potential biomarkers. These findings highlight the potential of PBMCs in the diagnosis and treatment of MDD, reinforcing the development of future neuroimmune-targeted therapies for depression.

## INTRODUCTION

Major depressive disorder (MDD) is a multifaceted psychiatric disorder^1^ characterized by persistent sadness, anhedonia, and a range of emotional and physical symptoms that significantly impair daily tasks^2^, making it a leading cause of disability worldwide^3^. The complexity of MDD has driven researchers to investigate its underlying mechanisms through multiple perspectives, including genetic, environmental, and neurobiological factors^4–6^. A growing body of evidence highlights the importance of neuroimmune crosstalk^7,8^, i.e., the bidirectional communication between the nervous and immune systems^9^, whereby inflammatory processes can influence neurobiological functions and vice-versa, contributing to the pathophysiology of depression^10,11^.

The neuroimmune interaction involves, among others, a network of soluble factors, including cytokines and neurotransmitters^12,13^, which can modulate neuronal activity and synaptic plasticity^12,14,15^. Building on these findings, we hypothesized that neuroimmune crosstalk extends beyond conventional paradigms. To explore this, we investigated the presence of an intrinsic network of neuronal molecules within peripheral blood mononuclear cells (PBMCs), aiming, through an integrative systems approach, to identify molecules that reflect those found in brain regions linked to depression. Integrative systems analysis is a multifaceted approach that combines high-dimensional data from diverse biological sources to elucidate complex interactions within biological systems^16^.

We employed RNA sequencing (RNA-seq) datasets to analyze gene expression profiles from PBMCs, and PBLs and several brain regions that can be associated with depression, including anterior cingulate cortex (ACC), anterior insula (aINS), cingulate gyrus 25 (Cg25), dorsolateral prefrontal cortex (DLPFC), nucleus accumbens (Nac), orbitofrontal cortex (OFC), and subiculum (Sub). This approach allowed the identification of shared molecular signatures that bridge the immune and nervous systems^17^. By meta-analysis, we unraveled a potential intricate network of gene interactions and biological processes (BPs) that underpin the neuroimmune crosstalk. This approach is crucial for understanding the systemic nature of depression, as it reveals how alterations in leukocyte gene expression may reflect or influence neurobiological changes in the brain. Furthermore, it enables us to identify potential biomarkers and therapeutic targets that can be explored for innovative treatment strategies.

## MATERIAL AND METHODS

### Data curation

Eligible studies were identified using PubMed, Google Scholar, and Gene Expression Omnibus (GEO) by searching for peer-reviewed datasets indexed with the terms “major depressive disorder,” “Homo sapiens,” and expression profiles from “high throughput sequencing” (bulk or single-cell RNA-seq), containing healthy controls and a minimum of 10 total samples. Initially, 58 studies reporting transcriptional data from human subjects were identified (**Supplementary Figure 1a**).

Studies were excluded based on the following criteria: (1) studies focused on other categories of depression or not identified as MDD datasets (n = 36) and other category of depression (n = 1); (2) studies not available in the databases, i.e., only found in original articles (n = 4); (3) studies that used single-nucleus RNA sequencing (snRNA-Seq) (n = 3); (4) datasets containing neurons, astrocytes, or immortalized B lymphocyte cell lines (n = 5); (5) datasets with fewer than 50 significant DEGs (n = 2); (6) datasets with fewer than 10 samples (n = 1); and (7) datasets associated with other psychiatric disorders, such as schizophrenia or bipolar disorder (n = 1). After applying these exclusion criteria, five datasets remained eligible in our workflow (**Suppl. Table S1a**), along with four datasets from the Wittenberg meta-analysis^18^ used for comparison with our meta-analysis. Together, these datasets included a total of 3,072 participants: 1,864 with MDD and 1,208 healthy controls (**Supplementary Figure 1b**).

These datasets were derived from 10 publications and were accessible for analysis: PBMCs from Trang et al.^19^ (https://github.com/insilico/DepressionGeneModules/tree/master/secondary_expression_data) and Cathomas et al. (GSE185855)^20^; ACC from Ramaker et al. (GSE80655)^21^ and Oh et al. (GSE193417)^22^; Labonté et al. (GSE102556)^23^; the study by Wittenberg et al.^24^ which includes the datasets by Leday et al. (GSK-HiTDIP cohort)^25^, Mostafavi et al.^26^, Jansen et al.^27^, Leday et al. (Janssen-BCR cohort)^28^, included here for comparison (**Supplementary Figure 1b**).

To comprehensively evaluate the transcriptional overlap between meta-differentially expressed genes (metaDEGs) (obtained through meta-analysis as described below) in PBMCs with DEGs from some brain regions, we used the dataset from Labonté et al.^29^ alongside the metaDEGs from ACC.

### Differential expression analysis

DEGs between groups were identified using DESeq2^30^. A statistical cutoff of adjusted p-value < 0.05, using FDR adjustment, was applied to define significant expression changes. For the Labonté et al.^29^ dataset, a p-value < 0.05 was used, consistent with the methodology of the original study. The analysis was conducted in R version 4.0.5. To address confounding variables (e.g., age, sex, and other available metadata parameters), was employed, ensuring that these factors did not confound the results. No significant effect of these factors on gene expression was observed beyond the primary influence of diagnosis (MDD vs. Control), affirming that the findings reflect genuine biological variations rather than technical artifacts.

### Meta-analysis of gene expression datasets

A comprehensive gene expression meta-analysis was conducted using the ‘MetaVolcanoR’ package^31^, applying the ‘Combining MetaVolcano’ approach. This analysis integrated datasets of PBMCs (Trang et al.^19^ and Cathomas et al.^32^) or ACCs (Ramaker et al. and Oh et al.^21^). Standard parameters were applied uniformly to ensure consistency across the results. The ACC region was selected for meta-analysis as it was the only brain region with more than two datasets that met the inclusion criteria, but also in consistency with findings from other studies, which highlight the fundamental role of this region in disorders such as MDD^33^. This approach combines DEGs from multiple bulk RNA-seq studies by calculating the mean and synthesizing gene differential expression using Fisher’s combined p-value method for each study (PBMC and ACC). The identified metaDEGs were further characterized and classified into gene types (e.g., protein-coding, miRNA, snoRNA, lncRNA, non-coding) using the ShinyGO^34^ online platform, providing a detailed understanding of the genomic landscape underlying the observed expression changes.

### Functional enrichment, clustering, and interactome

GO enrichment analysis was conducted using the EnrichR^35^ web tool to identify enriched BPs with a significance threshold of p-value < 0.05. To reduce redundancy in GO terms, we clustered similar terms based on semantic similarity, grouping BPs into parent categories to provide a more concise topological visualization of neuroimmunological BPs. This was achieved using the ‘rrvgo’^36^ package in R. For evaluating the nervous and immune clusters formed by BPs, the ‘appyters’^37^ web tool, integrated with EnrichR, was used. This approach allowed for identifying enriched BPs, organized into clusters on a scatter plot. The Leiden algorithm was applied for clustering, which reduced dimensions using uniform manifold approximation and projection (UMAP). Additionally, a network analysis of metaDEGs and BPs was performed to examine the neuroimmunological interactome, utilizing the ‘ggnet2’^38^ package in R.

### Characterization of synapse-related BPs

To characterize the metaDEGs associated with synaptic ontologies, we used the synaptic gene ontologies (SynGO)^39^ web tool through the EnrichR^35^ interface integrated within SynGO. This platform provides a robust framework for exploring gene functions specific to synapses, allowing us to gain deeper insights into the functional relationships between the identified metaDEGs and synaptic processes. By employing SynGO, we enhanced our understanding of how these genes from BPs and CC (cellular components) contribute to synaptic biology and their potential implications in the context of our study.

### Linear discriminant and principal component analyses of synapse-related genes

To assess the impact of nervous and immune genes in stratifying MDD and healthy individuals, we applied linear discriminant analysis (LDA)^40,41^ to PBMC datasets (Cathomas et al.^32^ and Trang et al.^19^, respectively), focusing on DEGs associated with synaptic ontologies (p < 0.05). LDA, a dimensionality reduction technique^42^, identifies the linear combinations of variables that maximize the separation between predefined groups (control vs. MDD).

Using a supervised learning approach, we trained the LDA model with 70% of the samples and tested it on the remaining 30%^42,43^. The model demonstrated over 60% accuracy in distinguishing control from MDD in both studies, identifying 49 common DEGs that effectively differentiated the groups. To further explore the stratification capacity of these 49 DEGs, we conducted PCA with spectral decomposition on both PBMC datasets. PCA was performed using the R functions prcomp, through factoextra package (PCA in R: prcomp)^44^.

### False discovery rate and binomial logistic regression of DEGs

We performed a Wilcoxon test using False Discovery Rate (FDR) on the 49 DEGs. Given the non-normal distribution of gene expression data, we used the Wilcoxon test to compare expression levels between diagnostic groups. P-values were adjusted with the FDR method, and boxplots were generated to visualize significant differences in gene expression, highlighting genes with adjusted p-values < 0.05 as significant. To further investigate the relationship between DEGs and the binary outcome of MDD versus control, we applied binomial logistic regression^45,46,47^ to the DEGs found significant in the Wilcoxon test. The logistic regression was implemented using the glm function from the base R package, with statistical significance set at p < 0.05.

### Construction of the “diseasome” network

To build the diseasome^48,49^ network mapping associations between genes and diseases based on shared genetic and molecular factors, we selected 11 DEGs common to the PBMC meta-analysis from LDA and seven brain regions (ACC meta-analysis, aINs, Cg25, DPLFC, OFC, Nac, and Sub). Using the EnrichR interface, we accessed the DisGeNET^49^ database to retrieve information on gene-disease associations. For the network visualization, we focused on diseases with a p-value < 0.05, enabling us to highlight the most statistically significant and relevant disease connections.

Additionally, we built a separate diseasome network for the DEGs significant after FDR adjustment, again using the EnrichR interface and focusing on gene-disease associations in DisGeNET with a p-value < 0.05 to emphasize the most impactful disease links.

### Tables employed to generate and comprehensively visualize the study’s results

The data/tables utilized to generate and visualize the results of our study are provided as supplementary files. These tables include comprehensive data inputs used for the analysis and detailed output results that encapsulate the findings and insights derived from our research. The supplementary input tables outline the dataset structure, variables, and relevant parameters, while the output tables display processed results, statistical metrics, and visual summaries that support our conclusions.

## RESULTS

### Intrinsic neuroimmunological networks in PBMCs modulated by MDD

To test our hypothesis about the presence of an intrinsic neuroimmune network within leukocytes^11,50,51^, we used PBMCs, a crucial component of the peripheral immune system. These cells offer a readily accessible source to test our hypothesis in MDD. Their role in bridging the immune and central nervous systems makes PBMCs^52^ ideal candidates for exploring neuroimmune interactions relevant to MDD pathology^53–55^.

Meta-analysis of the two bulk RNA-seq publically available across transcriptomic databases, using MetaVolcanoR, which combines differential gene expression results, revealed 903 down and 480 upregulated metaDEGs (p<0.05; **1a**), respectively. Most of these genes primarily code for proteins (**Figure 1b**). GO enrichment analysis of the metaDEGs revealed clusters of neuroimmunological BPs associated with immune and nervous system functions (**Figure 1c**). We identified 41 immune-related BPs (e.g., positive regulation of leukocyte chemotaxis and lymphocyte proliferation; **Figure 2**) significantly enriched by downregulated metaDEGs and 17 by upregulated metaDEGs. In contrast, downregulated metaDEGs linked to nervous system processes (e.g., synaptic vesicle recycling and neural tube development; **Figure 2**) significantly enriched ten BPs, while upregulated metaDEGs showed no significant enrichment (**Figures 1d**). Some metaDEGs belong to both nervous and immune systems, including genes like *OAS1*, *ISG5*, and *TRIM11*, which enrich the response to type I interferon (*IFN*; **Figure 2**), a BPs associated with the development of MDD^26^.

**Figure 1.**
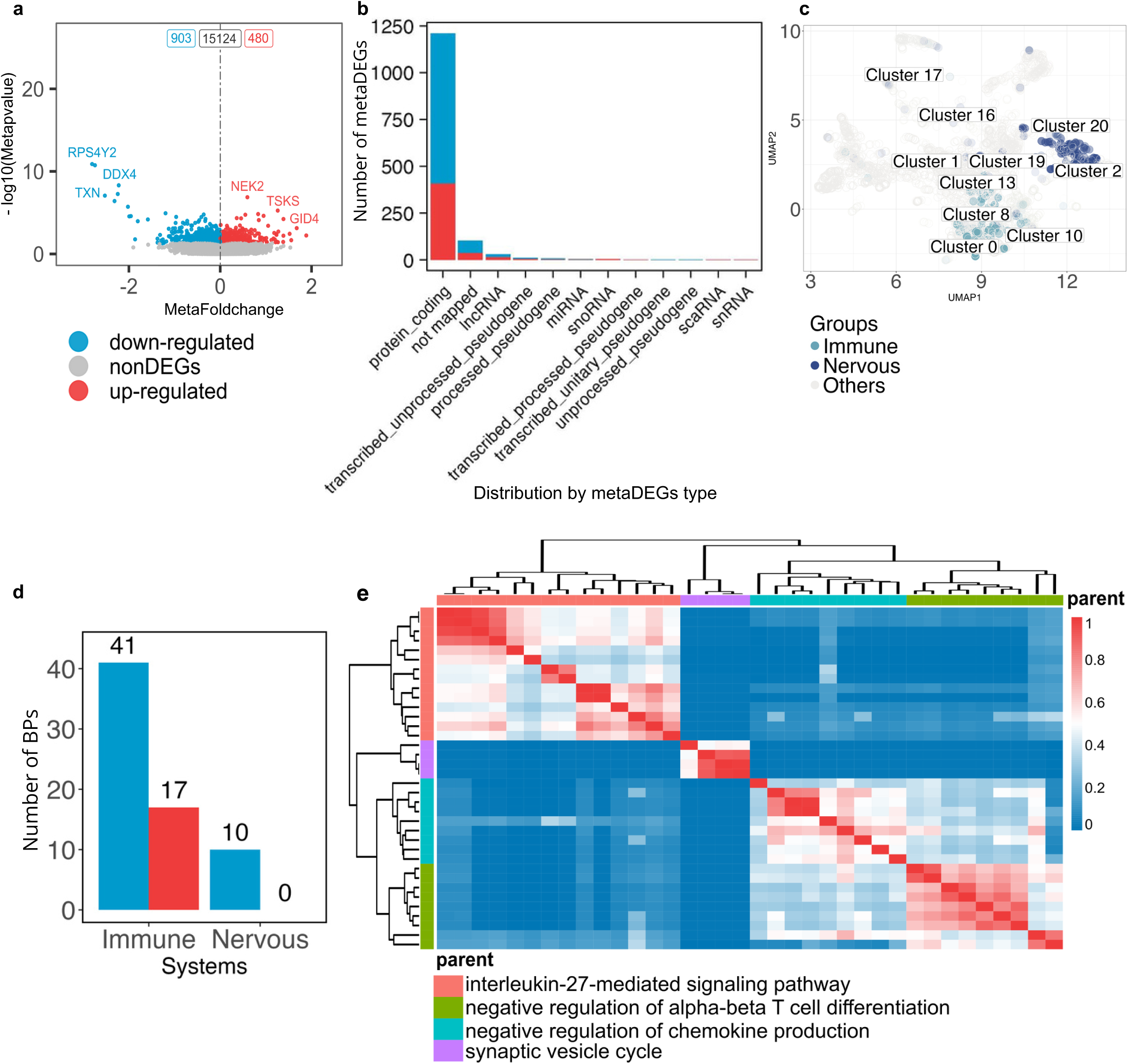
Meta-analysis of PBMCs and enrichment of biological processes (BPs). (a) Volcano plot displaying down- and up-regulated metaDEGs. (b) Bar plot of up-regulated (red) and down-regulated (blue) metaDEGs gene types. (c) Scatterplot of enriched BPs related to immune and nervous systems, with color intensity representing statistical significance. (d) Bar plot showing immune- and nervous system-related BPs. (e) Heatmap of the similarity matrix for BPs, as analyzed using rrvgo, demonstrating clustering of related processes.

**Figure 2.**
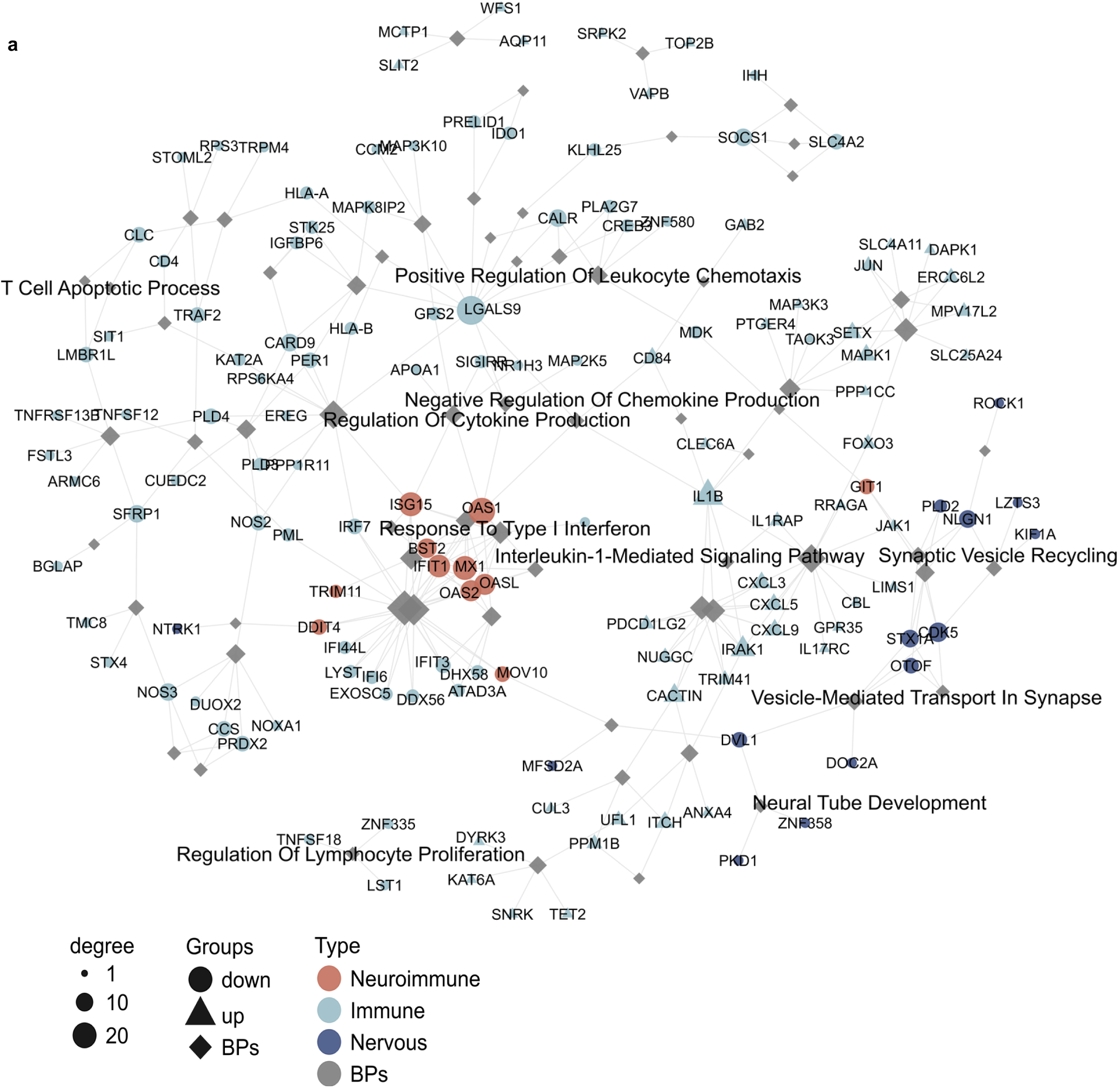
Network of immune- and nervous system-related biological processes (BPs). (a) Network visualization showing the relationships among BPs associated with immune and nervous systems. MetaDEGs that participate in both immune and nervous processes are labeled as neuroimmune DEGs and are highlighted in light red, indicating their dual role in neuroimmune interactions.

Moreover, we utilized GO annotation to interpret the biological relevance of DEGs within PBMCs, leveraging the rrvgo^56^ package to reduce redundancy in enriched GO terms and facilitate interpretation. In addition to identifying dysregulation in immune-related BPs, such as granulocyte activation, our results also highlighted the presence of nervous system-related BPs within PBMCs as parent terms in the GO hierarchy (**Figure 1e**). These included vital processes such as synaptic vesicle cycle.

The detection of these nervous system-related BPs within immune cells underscores the potential role of PBMCs in the neuroimmune crosstalk involved in MDD^53–55^, further supporting the hypothesis of leukocytes contributing to both immune and neural regulation.

### Synaptic-related BPs present in PBMCs

To deepen our understanding of the involvement of nervous system-related BPs, we focused on characterizing the presence of neurotransmitter secretion and other synaptic-related BPs in PBMCs. To achieve this, we used the SynGO^39^ tool, an advanced resource that facilitates the systematic annotation of synaptic DEGs. SynGO provides an interactive knowledge base and ontology for synaptic locations and processes based on expert-curated evidence. This tool enabled us to explore how synaptic dysregulation, a key feature in several brain disorders (termed “synaptopathies”)^57^, is reflected in our PBMC data.

SynGO annotation identified that 117 out of the 1,383 metaDEGs in PBMCs (75 downregulated and 42 upregulated) corresponded to synapse-related genes (**Figure 3a**). This finding suggests that PBMCs from patients with MDD express genes typically associated with synaptic functions, underscoring the potential for neuroimmune crosstalk within these immune cells. Among the top five synaptic ontologies enriched by upregulated metaDEGs in PBMCs are critical processes such as the regulation of synaptic vesicle exocytosis, postsynaptic density, and presynaptic functions (**Figure 3b**). Similarly, downregulated metaDEGs also enriched synaptic-related BPs, including regulating postsynaptic assembly, anchored presynaptic membrane components, and regulating neurotransmitter receptor localization to the postsynaptic specialization membrane.

**Figure 3.**
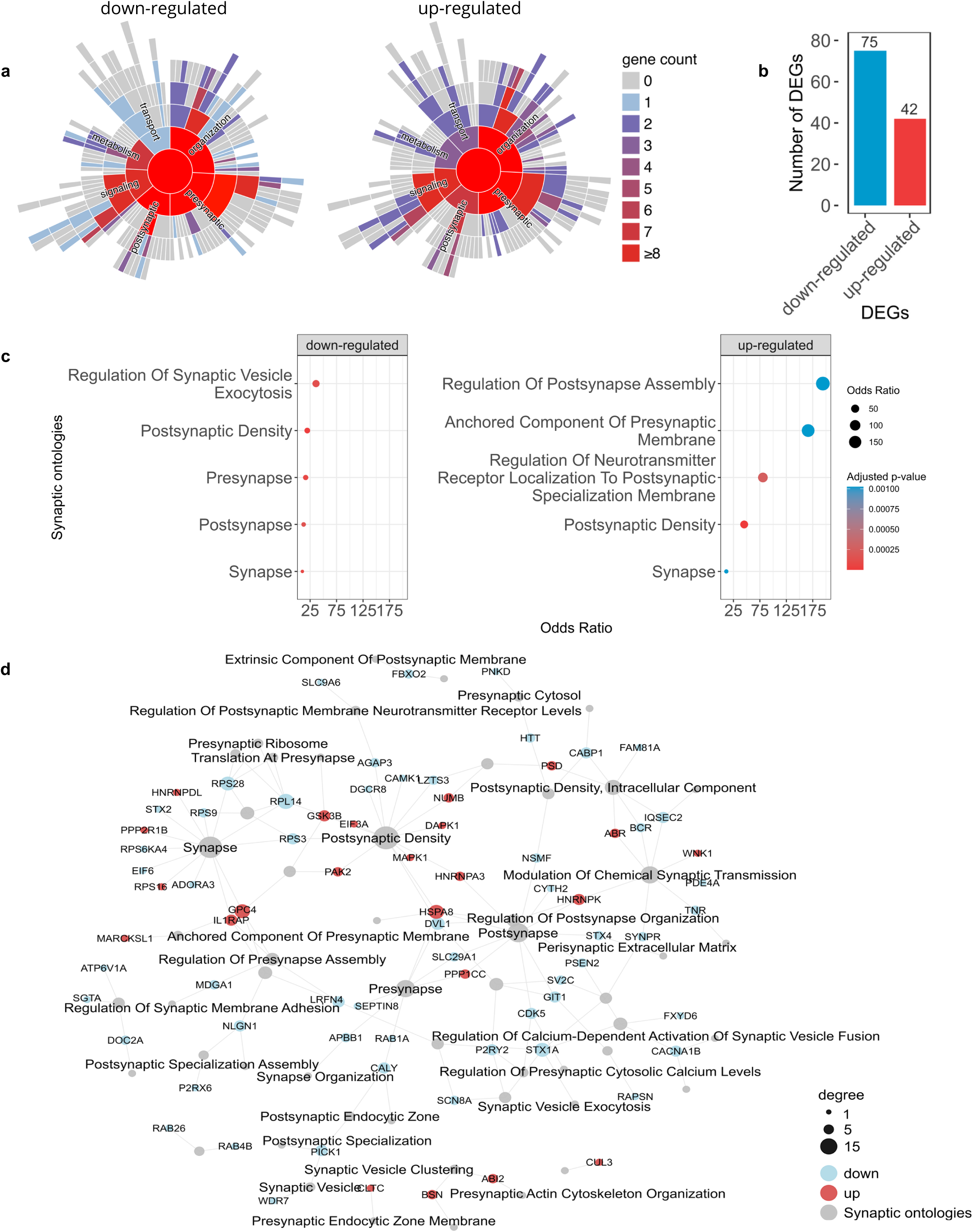
Synapse-related ontologies from PBMC meta-analysis. (a) Synaptic ontologies identified via SynGO, with each radial segment representing gene counts associated with specific synaptic functions. (b) Bar plot showing the distribution of metaDEGs that are enriched in synaptic ontologies. (c) Re-enrichment analysis of synapse-related metaDEGs, highlighting the five most significant synaptic ontologies for both up- and down-regulated genes. (d) Network illustrating synapse-associated metaDEGs, with a comparative view of up- (red) and down-regulated metaDEGs (blue).

Notably, the SynGO analysis of the metaDEGs identified in the Wittenberg et al.^18^ study revealed several synaptic-related genes, including five that overlap with the metaDEGs obtained from our PBMC meta-analysis (**Supplementary Figure 2**). This discovery underscores the potential role of synaptic gene dysregulation within the molecular landscape of the condition being studied, supporting the hypothesis that synaptic-related alterations in PBMCs may be central to its pathology.

Network analysis revealed complex interactions among synapse-related genes and ontologies, emphasizing the interconnected nature of the neuroimmune network within PBMCs (**Figure 3c**).

### Synaptic-related genes in PBMCs differentiate MDD patients from healthy controls

Investigating synaptic-related genes in PBMCs could provide critical insights into neuroimmune interactions, opening pathways for innovative diagnostic and therapeutic approaches for MDD. To pursue this, we applied stratification techniques, specifically LDA and PCA, after identifying the most relevant genes. This selection was based on an enrichment analysis using the SynGO tool via EnrichR of 117 synaptic-related metaDEGs and the intersection of shared genes between the Cathomas et al. and Trang et al. datasets (**Figure 4a**).

**Figure 4.**
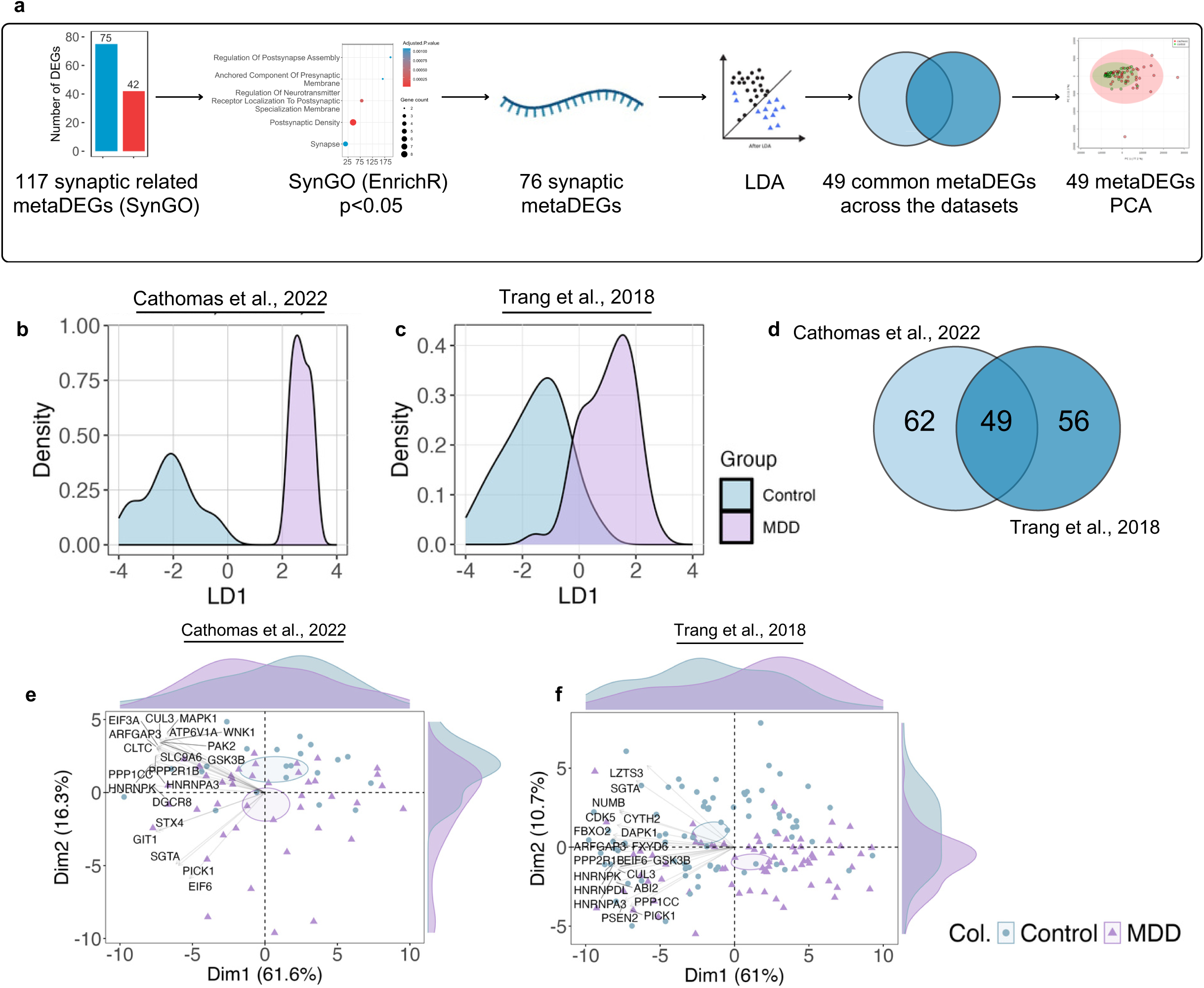
Ranking and stratification of synaptic-related metaDEGs. (a) Flowchart summarizing the steps of Linear Discriminant Analysis (LDA) used for ranking and stratifying metaDEGs. (b-c) Density plots of the first Linear Discriminant (LD1) axis, showing the separation between metaDEGs in healthy controls (blue) and MDD (purple) groups for datasets from Cathomas et al. (2022) and Trang et al. (2018), respectively. (d) Venn diagram showing the overlap of metaDEGs identified by LDA across the two datasets, with 62 DEGs in Cathomas et al. and 56 in Trang et al. (e-f) Principal Component Analysis (PCA) plots displaying the distribution of the 49 overlapping metaDEGs along PC1 and PC2 axes, highlighting the separation between MDD and healthy controls from Cathomas et al. (left) and Trang et al. (rigth) datasets.

LDA analysis identified 76 metaDEGs associated with enriched BPs that effectively distinguished MDD patients from healthy controls (**Figures 4b, c**). Among these, 56 DEGs were present in the Trang et al. dataset and 62 in the Cathomas et al. dataset. Notably, 49 synaptic-related genes were common to both datasets (**Figure 4d**), serving as robust markers for differentiating MDD patients from healthy controls in PBMC datasets, as demonstrated by PCA (**Figures 4e, f**).

### Synaptic-related genes shared between PBMCs and brain regions in MDD

The findings above indicate the involvement of synaptic-related genes in PBMCs, suggesting that immune cells might reflect specific neural characteristics associated with MDD^6^. To investigate this possibility, we conducted a consensus analysis of 49 common synaptic-related metaDEGs, assessing their expression across seven brain regions (**Figure 5a**). These included the ACC from our meta-analysis (**Supplementary Figure 3**) and six additional areas from the Labonté et al. (2017) dataset: aINS, Cg25, DLPFC, Nac, OFC, and Sub.

**Figure 5.**
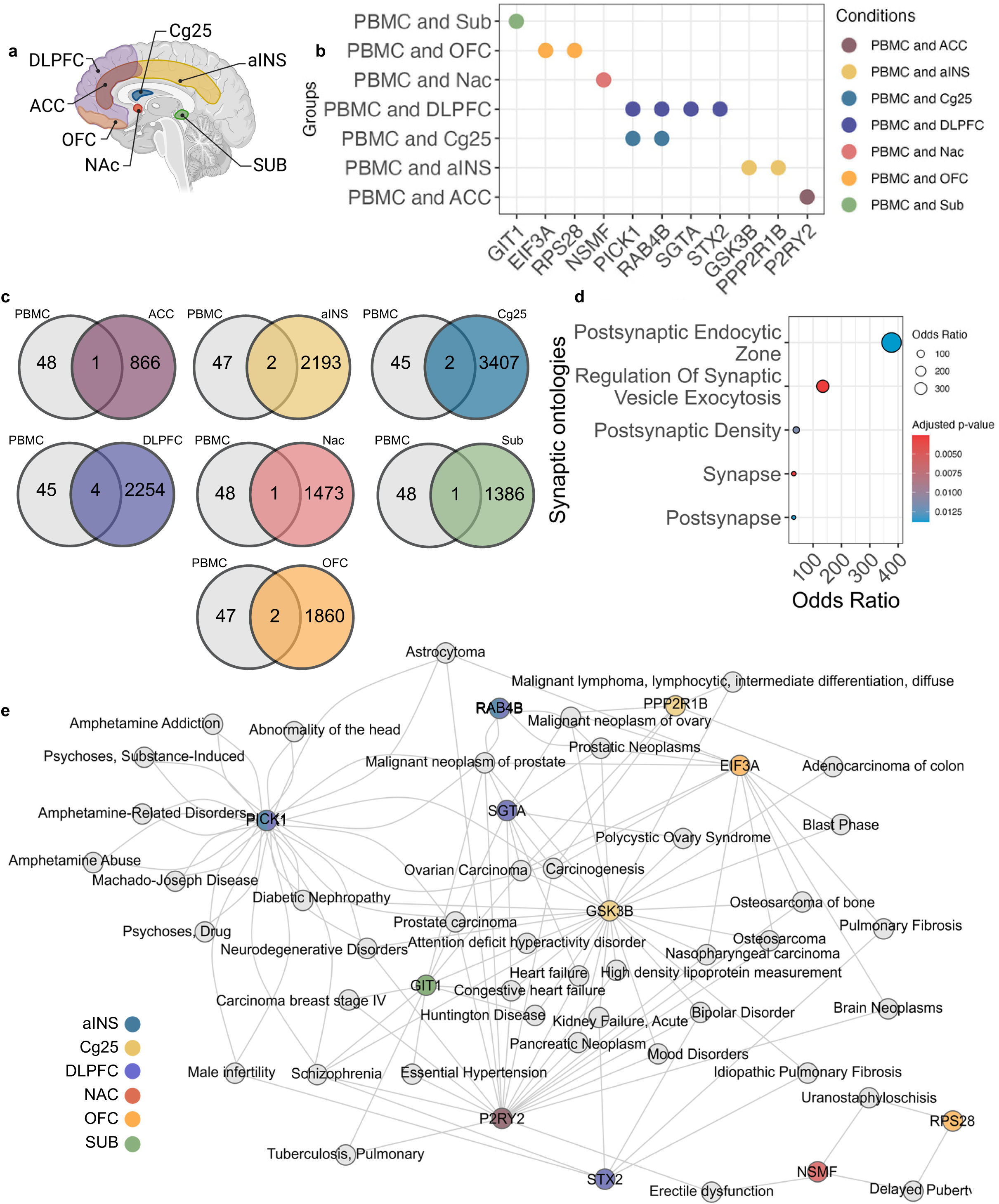
Intersection of DEGs from PBMC meta-analysis and brain regions in MDD. (a) Anatomical locations of the brain regions analyzed in the study, including the Anterior Cingulate Cortex (ACC), Anterior Insula (aINS), Cingulate Gyrus Area 25 (Cg25), Dorsolateral Prefrontal Cortex (DLPFC), Nucleus Accumbens (Nac), Orbitofrontal Cortex (OFC), and Subiculum (Sub). (b) Dot plot illustrating the 11 DEGs that are common to both PBMCs and each brain region. (c) Venn diagram depicting the overlap between the 49 common DEGs identified in PBMCs (Figure 5d) and the total DEGs from the brain regions. (d) Bar plot displaying the top 5 significant synaptic ontology enrichments derived from the 11 common DEGs. (e) Disease network analysis of the 11 common DEGs shared between PBMCs and brain regions, showing only significant (p < 0.05) gene-disease associations.

This approach identified 11 synaptic-related genes shared between PBMCs and these brain regions (**Figures 5b, c**) GIT1^6,58^ EIF3A^59,60^, RPS28^60–62^, NSMF^63–65^, PICK1^66,67^, RAB4B^68^, SGTA^68,69^, STX2^70,71^, GSK3B^72–74^, PPP2R1B^75,76^, and P2RY2^77–79^. Their association with synaptic processes is illustrated in **Figure 5d**.

Further diseasome analysis mapped these DEGs to a spectrum of health conditions, including psychoneurological disorders (e.g., mood disorders^74,80,81^, bipolar disorder^81,82^, neurodegenerative diseases^83–85^, psychoses^86^, schizophrenia^86–89^, ADHD ^90,91^, Huntington’s disease ^92,93^), metabolic diseases (e.g., diabetic nephropathy^94,95^, HDL abnormalities^96,97^), cardiovascular diseases (e.g., heart failure^69,98,99^, hypertension^100,101^), pulmonary diseases (e.g., pulmonary fibrosis^102,103^, tuberculosis^104,105^), cancer (e.g., brain neoplasm^106–108^, astrocytoma^106,109,110^, lymphoma^111,112^, ovarian^113–116^ pancreatic cancer^117,118^, osteosarcoma^119,120^), and reproductive disorders (e.g., male infertility^121,122^, erectile dysfunction^123,124^, polycystic ovarian syndrome^125,126^) (**Figure 5e**).

### Synaptic-related gene predictors in MDD

To uncover the most robust synaptic-related genes that might predict MDD, we applied a Wilcoxon analysis using FDR method to narrow down the pool of 49 synaptic-related genes to those with the most statistically significant associations. We then used binomial regression to further validate these associations. This two-step approach helped us pinpoint five genes (*ADORA3*^126,127^, *FBXO2*^127,128^, *FXYD6*^129^, *RPS28*^61^, and *RPS9*^61^) that remained statistically significant (**Figure 6a**) and demonstrated a strong predictive relationship with MDD based on the regression analysis (**Figure 6b**).

**Figure 6.**
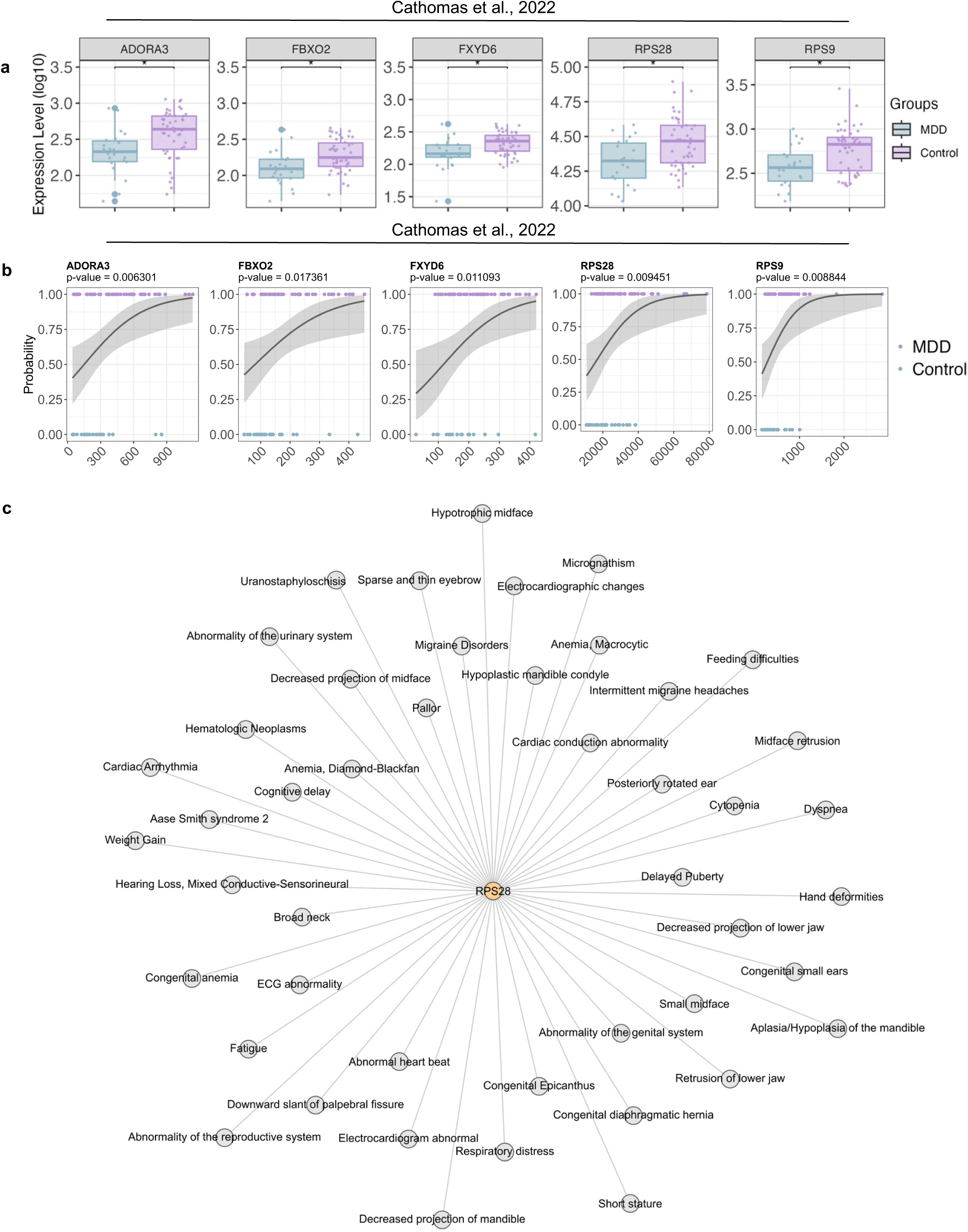
Synaptic-related DEGs which are FDR significant and binomial logistic regression analysis. (a) Boxplot illustrating the five synaptic metaDEGs synaptic-related that were significant after false discovery rate (FDR) adjustment. The study by Cathomas et al. (2022) identified the following significant DEGs: ADORA3, FBXO2, FXYD6, RPS28, and RPS9. No FDR-significant DEGs were found in the study by Trang et al. (2018). (b) Scatterplots showing the results of binomial logistic regression for the expression of each metaDEG synaptic-related, comparing the MDD group to healthy controls based on data from Cathomas et al. (2022). (c) Diseasome network focusing on RPS28, showing only significant (p < 0.05) gene-disease associations.

While four of these genes were dysregulated only in PBMCs, *RPS28* exhibited dysregulation in both PBMCs and a brain region tied explicitly to mood and cognition, the orbitofrontal cortex (OFC) (**Figure 5**). Further exploration through diseasome analysis revealed that *RPS28* is associated with a range of symptoms often seen in MDD. In addition to its role in, *RPS28* has connections to symptoms like weight gain^130^, fatigue^130,131^, intermittent migraine headaches^132,133^, and cognitive delay^134,135^ (**Figure 6c)**.

## DISCUSSION

Our study uncovers an intricate neuroimmune network within PBMCs that is disrupted in MDD. This discovery shifts the traditional understanding of the neuroimmune interface (crosstalk)^53,55,136^ by suggesting that immune cells harbor synaptic machinery and are thus not passive players but active intrinsic contributors to neurobiological processes relevant to mood disorders^137^. Identifying specific synaptic-related genes within PBMCs provides compelling evidence of molecular similarities between leukocytes and brain regions critically associated with MDD. This finding reinforces that PBMCs may participate in a feedback loop with neural circuits^138,139^, potentially affecting mood regulation^139–141^. The capacity of PBMCs to synthesize neurotransmitters^142,143^ (e.g., serotonin^143^, dopamine^143,144^) indicates a self-modulatory function, also through an autocrine manner^143,145^, that could influence their implications in health and disease states.

The identification of 117 synaptic-related metaDEGs in PBMCs from MDD patients, involving processes such as synaptic vesicle exocytosis and postsynaptic assembly, suggests a systemic role of the neuroimmune axis in MDD. Notably, the capacity of 49 of these metaDEGs to distinguish MDD patients from healthy controls underscores their potential as robust peripheral markers, reflecting key aspects of neuroimmune dysregulation associated with MDD. Through LDA and PCA, we observed a separation between MDD and control groups, highlighting the promise of these PBMC-derived genes, alongside previously reported immune-related markers (e.g. C-reactive protein^146^, *IL-12*^147^, *P2X7* (Drevets et al., 2022) and *PAX6*^148^), as accessible indicators of neurobiological alterations linked to MDD.

Among the 49-stratifying synaptic-related genes, 11 DEGs were shared between PBMCs and brain regions associated with mood regulation. Genes like *GIT1*, *PICK1*, and RAB4B are integral to synaptic processes^58,66–68^ such as vesicle transport^149–151^ and receptor trafficking^152,153^, which are critical for neural connectivity and cognitive functions^154,155^. Their presence in both immune and neural compartments supports the hypothesis that PBMCs mirror molecular processes in the brain and may play an active role in neuroimmune communication.

Further, the diseasome network analysis revealed associations of these 11 shared genes with various psychiatric and physiological conditions, including mood disorders^74,80,81^, neurodegenerative diseases^83,85,156^, and cardiovascular issues^95,100,101,157^. For instance, *PICK1’s* involvement in *AMPA* receptor trafficking^152^, essential for synaptic transmission^152^ and plasticity^14,152,158^, directly relates to MDD symptoms like anhedonia^159,160^ and cognitive impairment^83,160,161^. Similarly, *RAB4B*, which regulates endocytic recycling^151^, may influence signaling pathways relevant to both immune responses and neural communication. These connections suggest that these genes may serve as potential therapeutic targets for conditions beyond MDD, offering insights into the broader systemic impact of MDD on physical health.

The diseasome network analysis aligns with the observed multimorbidity in MDD^162^ patients, highlighting the intricate interconnections between mental and physical health^163^. These findings underscore how inflammatory, neuroimmune, and neurodegenerative mechanisms^164^ contribute to association of MDD with multiple chronic conditions, such as cardiovascular disease^165^, metabolic syndrome^166^, autoimmune disorders^167^, and neurodegenerative diseases^168^. For instance, genes like *RAB4B* and *ADORA3*, which play roles in immune response^169,170^ and vascular regulation^171,172^, may explain the coexistence of chronic inflammation in both MDD and cardiovascular diseases^165^, where elevated pro-inflammatory cytokines can impair endothelial function, exacerbate oxidative stress, and increase cardiovascular risk^173,174^. Similarly, *PICK1* and *RPS28*, both involved in synaptic plasticity^175^, connect MDD with neurodegenerative conditions^168^, suggesting that chronic neuroinflammation in depression may accelerate neuronal loss and cognitive decline^176^. Recognizing these overlapping pathways points to the need for integrated treatment approaches that address both the symptoms of MDD and its systemic effects, offering potential for interventions that reduce inflammation, support neuroprotection, and improve cardiovascular and metabolic health, ultimately addressing the broad multimorbidity associated with depression.

Of note, five synaptic-related DEGs, *ADORA3, FBXO2, FXYD6, RPS28*, and *RPS9*, remained significant after FDR adjustment, underscoring their robust association with MDD. These genes play diverse roles in synaptic and immune functions, linking them to MDD pathology and broader physiological processes. *ADORA3*, an adenosine receptor, has immunomodulatory effects^120^ that may contribute to the inflammatory profile observed in MDD patients. Dysregulation in the adenosine pathway can impact neurotransmission^177^ and neuronal plasticity^14^, indicating that *ADORA3* might influence both immune and mood-regulating functions^80^.

*RPS28,* a ribosomal protein identified as dysregulated in both PBMCs and the OFC, emerges as a particularly promising biomarker due to its established links with cognitive and mood disorders. Associated with symptoms such as cognitive delay^134,135^ and fatigue^131,178^, *RPS28* could be integral to understanding the systemic manifestations of MDD. The diseasome network for *RPS28* highlights its association with metabolic^179^, cardiovascular^157^, and neuropsychiatric disorders^132,134,180^, further supporting a model where MDD extends beyond mental health to impact physical health^181^ via systemic pathways.

In conclusion, these findings provide a foundation for innovative diagnostic tools and treatments that integrate the neuroimmune axis. The 49-stratifying synaptic-related genes, especially the 11 DEGs shared between PBMCs and brain regions, offer a molecular fingerprint that could be used for peripheral diagnostics. Moreover, the five FDR-significant DEGs present therapeutic potential, paving the way for interventions targeting the neuroimmune dysregulation central to MDD.

Future research should validate these findings across larger cohorts and investigate whether modulating these gene expressions in PBMCs affects mood-related behaviors. Given the overlapping roles of these genes in immune and neural functions, therapeutic approaches could be developed to address MDD neuroimmune and neurobiological dimensions, potentially providing relief for patients who do not respond to conventional treatments.

## Supporting information

Supplementary Informations

## Acknowledgements

We thank the São Paulo Research Foundation (FAPESP grants 2018/18886-9 to OCM, 2023/12268-0 to ASA, 2023/13356-0 to DLMF, 2023/07806-2 to ISF, 2023/06086-6 to PMB) for financial support. We acknowledge the National Council for Scientific and Technological Development (CNPq) (grants: 309482/2022-4 to OCM and 102430/2022-5 to LFS, 130027/2023-5 to YLGC), Foundation Coordination for the Improvement of Higher Education Personnel (CAPES) (grants: 88887.801068/2023-00 ALN, 88887.992612/2024-00 to IAFB, 88887.699840/2022-00 to FYNV) and Serrapilheira fellowship (grant: #R2111-39828 to DGA).

## References

1. Kim, Y.-K. Major Depressive Disorder: Current Advances and Paradigm Shifts. Psychiatry Investig 17, 179–180 (2020).

2. Cui, L. et al. Major depressive disorder: hypothesis, mechanism, prevention and treatment. Signal Transduct Target Ther 9, 30 (2024).

3. James, S. L. et al. Global, regional, and national incidence, prevalence, and years lived with disability for 354 diseases and injuries for 195 countries and territories, 1990–2017: a systematic analysis for the Global Burden of Disease Study 2017. The Lancet 392, 1789–1858 (2018).

4. aan het Rot, M., Mathew, S. J. & Charney, D. S. Neurobiological mechanisms in major depressive disorder. Can Med Assoc J 180, 305–313 (2009).

5. Dean, J. & Keshavan, M. The neurobiology of depression: An integrated view. Asian J Psychiatr 27, 101–111 (2017).

6. Fries, G. R., Saldana, V. A., Finnstein, J. & Rein, T. Molecular pathways of major depressive disorder converge on the synapse. Mol Psychiatry 28, 284–297 (2023).

7. Wang, C., Zhou, Y. & Feinstein, A. Neuro-immune crosstalk in depressive symptoms of multiple sclerosis. Neurobiol Dis 177, 106005 (2023).

8. Mandolesi, G., Musella, A. & Centonze, D. Special Issue Title: The neuroimmune crosstalk in mood disturbances of neurological and psychiatric diseases. Neurobiol Dis 183, 106180 (2023).

9. Kamimura, D., Tanaka, Y., Hasebe, R. & Murakami, M. Bidirectional communication between neural and immune systems. Int Immunol 32, 693–701 (2020).

10. Beurel, E., Toups, M. & Nemeroff, C. B. The Bidirectional Relationship of Depression and Inflammation: Double Trouble. Neuron 107, 234–256 (2020).

11. Hodo, T. W., de Aquino, M. T. P., Shimamoto, A. & Shanker, A. Critical Neurotransmitters in the Neuroimmune Network. Front Immunol 11, (2020).

12. Dantzer, R. Neuroimmune Interactions: From the Brain to the Immune System and Vice Versa. Physiol Rev 98, 477–504 (2018).

13. Dantzer, R. Neuroimmune Interactions: From the Brain to the Immune System and Vice Versa. Physiol Rev 98, 477–504 (2018).

14. Ren, F. & Guo, R. Synaptic Microenvironment in Depressive Disorder: Insights from Synaptic Plasticity. Neuropsychiatr Dis Treat Volume 17, 157–165 (2021).

15. Kerage, D., Sloan, E. K., Mattarollo, S. R. & McCombe, P. A. Interaction of neurotransmitters and neurochemicals with lymphocytes. J Neuroimmunol 332, 99–111 (2019).

16. Greene, C. S. & Troyanskaya, O. G. Integrative Systems Biology for Data-Driven Knowledge Discovery. Semin Nephrol 30, 443–454 (2010).

17. Rosati, D. et al. Differential gene expression analysis pipelines and bioinformatic tools for the identification of specific biomarkers: A review. Comput Struct Biotechnol J 23, 1154–1168 (2024).

18. Drevets, W. C., Wittenberg, G. M., Bullmore, E. T. & Manji, H. K. Immune targets for therapeutic development in depression: towards precision medicine. Nat Rev Drug Discov 21, 224–244 (2022).

19. Le, T. T. et al. Correction: Identification and replication of RNA-Seq gene network modules associated with depression severity. Transl Psychiatry 10, 282 (2020).

20. Cathomas, F. et al. Whole blood transcriptional signatures associated with rapid antidepressant response to ketamine in patients with treatment resistant depression. Transl Psychiatry 12, 12 (2022).

21. Ramaker, R. C. et al. Post-mortem molecular profiling of three psychiatric disorders. Genome Med 9, 72 (2017).

22. Oh, H., Newton, D., Lewis, D. & Sibille, E. Lower Levels of GABAergic Function Markers in Corticotropin-Releasing Hormone-Expressing Neurons in the sgACC of Human Subjects With Depression. Front Psychiatry 13, (2022).

23. Mansouri, S. et al. Transcriptional dissection of symptomatic profiles across the brain of men and women with depression. Nat Commun 14, 6835 (2023).

24. Wittenberg, G. M., Greene, J., Vértes, P. E., Drevets, W. C. & Bullmore, E. T. Major Depressive Disorder Is Associated With Differential Expression of Innate Immune and Neutrophil-Related Gene Networks in Peripheral Blood: A Quantitative Review of Whole-Genome Transcriptional Data From Case-Control Studies. Biol Psychiatry 88, 625–637 (2020).

25. Leday, G. G. R. et al. Replicable and Coupled Changes in Innate and Adaptive Immune Gene Expression in Two Case-Control Studies of Blood Microarrays in Major Depressive Disorder. Biol Psychiatry 83, 70–80 (2018).

26. Mostafavi, S. et al. Type I interferon signaling genes in recurrent major depression: increased expression detected by whole-blood RNA sequencing. Mol Psychiatry 19, 1267–1274 (2014).

27. Jansen, R. et al. Gene expression in major depressive disorder. Mol Psychiatry 21, 339–347 (2016).

28. Leday, G. G. R. et al. Replicable and Coupled Changes in Innate and Adaptive Immune Gene Expression in Two Case-Control Studies of Blood Microarrays in Major Depressive Disorder. Biol Psychiatry 83, 70–80 (2018).

29. Mansouri, S. et al. Transcriptional dissection of symptomatic profiles across the brain of men and women with depression. Nat Commun 14, 6835 (2023).

30. Love, M. I., Huber, W. & Anders, S. Moderated estimation of fold change and dispersion for RNA-seq data with DESeq2. Genome Biol 15, 550 (2014).

31. Prada C, L. D. N. H. MetaVolcanoR: Gene Expression Meta-analysis Visualization Tool. . Bioconductor. R package version 1.16.0. (2024).

32. Cathomas, F. et al. Whole blood transcriptional signatures associated with rapid antidepressant response to ketamine in patients with treatment resistant depression. Transl Psychiatry 12, 12 (2022).

33. Zhong, X. et al. Integrated analysis of transcriptional changes in major depressive disorder: Insights from blood and anterior cingulate cortex. Heliyon 10, e28960 (2024).

34. Ge, S. X., Jung, D. & Yao, R. ShinyGO: a graphical gene-set enrichment tool for animals and plants. Bioinformatics 36, 2628–2629 (2020).

35. Chen, E. Y. et al. Enrichr: interactive and collaborative HTML5 gene list enrichment analysis tool. BMC Bioinformatics 14, 128 (2013).

36. Sayols, S. rrvgo: a Bioconductor package for interpreting lists of Gene Ontology terms. MicroPubl Biol 2023, (2023).

37. Clarke, D. J. B. et al. Appyters: Turning Jupyter Notebooks into data-driven web apps. Patterns 2, 100213 (2021).

38. Tyner, S., Briatte, F. & Hofmann, H. Network Visualization with ggplot2. R J 9, 27 (2017).

39. Koopmans, F. et al. SynGO: An Evidence-Based, Expert-Curated Knowledge Base for the Synapse. Neuron 103, 217–234.e4 (2019).

40. Edwin Munene Kagereki. Principal Component Analysis and Linear Discriminant Analysis in Gene Expression Data. (University of Nairobi Institute of Tropical and Infectious Diseases, Nairobi, 2013).

41. Wu, M. C., Zhang, L., Wang, Z., Christiani, D. C. & Lin, X. Sparse linear discriminant analysis for simultaneous testing for the significance of a gene set/pathway and gene selection. Bioinformatics 25, 1145–1151 (2009).

42. Sharma, A. & Paliwal, K. K. Cancer classification by gradient LDA technique using microarray gene expression data. Data Knowl Eng 66, 338–347 (2008).

43. Liu, H., Zeng, Y., Zhao, X. & Tong, H. Improved geographical origin discrimination for tea using ICP□MS and ICP□OES techniques in combination with chemometric approach. J Sci Food Agric 100, 3507–3516 (2020).

44. Kotlyar, M. et al. IID 2021: towards context-specific protein interaction analyses by increased coverage, enhanced annotation and enrichment analysis. Nucleic Acids Res 50, D640–D647 (2022).

45. Sarah Schwartz & Tyson Barrett. Encyclopedia of Quantitative Methods in R: 10. Logistic Regression - Ex: Depression (Hoffman). vol. 4 (2021).

46. Hoffman, J. I. E. Logistic Regression. in Biostatistics for Medical and Biomedical Practitioners 601–611 (Elsevier, 2015). doi:10.1016/B978-0-12-802387-7.00033-0.

47. Wickham, H. Ggplot2. (Springer International Publishing, Cham, 2016). doi:10.1007/978-3-319-24277-4.

48. Wysocki, K. & Ritter, L. Diseasome. Annu Rev Nurs Res 29, 55–72 (2011).

49. Loscalzo, J., Kohane, I. & Barabasi, A. Human disease classification in the postgenomic era: A complex systems approach to human pathobiology. Mol Syst Biol 3, (2007).

50. Kanashiro, A. et al. The role of neutrophils in neuro-immune modulation. Pharmacol Res 151, 104580 (2020).

51. Carson, M. J., Cameron Thrash, J. & Walter, B. The cellular response in neuroinflammation: The role of leukocytes, microglia and astrocytes in neuronal death and survival. Clin Neurosci Res 6, 237–245 (2006).

52. Sun, L. et al. Peripheral Blood Mononuclear Cell Biomarkers for Major Depressive Disorder: A Transcriptomic Approach. Depress Anxiety 2024, (2024).

53. Sarno, E., Moeser, A. J. & Robison, A. J. Neuroimmunology of depression. in 259–292 (2021). doi:10.1016/bs.apha.2021.03.004.

54. Abelaira, H. M., Silva, R. H., Carlessi, A. S., Quevedo, J. & Réus, G. Z. Neuro-Immune Interactions in Depression: Mechanisms and Translational Implications. in Neurobiology of Depression 75–88 (Elsevier, 2019). doi:10.1016/B978-0-12-813333-0.00008-1.

55. Hodes, G. E., Kana, V., Menard, C., Merad, M. & Russo, S. J. Neuroimmune mechanisms of depression. Nat Neurosci 18, 1386–1393 (2015).

56. Sayols, S. rrvgo: a Bioconductor package for interpreting lists of Gene Ontology terms. MicroPubl Biol 2023, (2023).

57. Lepeta, K. et al. Synaptopathies: synaptic dysfunction in neurological disorders – A review from students to students. J Neurochem 138, 785–805 (2016).

58. Martyn, A. C. et al. GIT1 regulates synaptic structural plasticity underlying learning. PLoS One 13, e0194350 (2018).

59. Liu, H.-H. et al. Role of the visual experience-dependent nascent proteome in neuronal plasticity. Elife 7, (2018).

60. Aguilar-Valles, A. et al. Translational control of depression-like behavior via phosphorylation of eukaryotic translation initiation factor 4E. Nat Commun 9, 2459 (2018).

61. Fusco, C. M. et al. Neuronal ribosomes exhibit dynamic and context-dependent exchange of ribosomal proteins. Nat Commun 12, 6127 (2021).

62. Sharma, V., Swaminathan, K. & Shukla, R. The Ribosome Hypothesis: Decoding Mood Disorder Complexity. Int J Mol Sci 25, 2815 (2024).

63. Spilker, C., Grochowska, K. M. & Kreutz, M. R. What do we learn from the murine *Jacob/Nsmf* gene knockout for human disease? Rare Diseases 4, e1241361 (2016).

64. Lau, C. G. & Zukin, R. S. NMDA receptor trafficking in synaptic plasticity and neuropsychiatric disorders. Nat Rev Neurosci 8, 413–426 (2007).

65. Amidfar, M. et al. The role of NMDA receptor in neurobiology and treatment of major depressive disorder: Evidence from translational research. Prog Neuropsychopharmacol Biol Psychiatry 94, 109668 (2019).

66. Perroy, J. PICK1 is required for the control of synaptic transmission by the metabotropic glutamate receptor 7. EMBO J 21, 2990–2999 (2002).

67. Xu, J., Kam, C., Luo, J. & Xia, J. PICK1 Mediates Synaptic Recruitment of AMPA Receptors at Neurexin-Induced Postsynaptic Sites. The Journal of Neuroscience 34, 15415–15424 (2014).

68. Kang, H. J. et al. Decreased expression of synapse-related genes and loss of synapses in major depressive disorder. Nat Med 18, 1413–1417 (2012).

69. Philp, L. K. et al. Small Glutamine-Rich Tetratricopeptide Repeat-Containing Protein Alpha (SGTA) Ablation Limits Offspring Viability and Growth in Mice. Sci Rep 6, 28950 (2016).

70. Salazar Lázaro, A., Trimbuch, T., Vardar, G. & Rosenmund, C. The stability of the primed pool of synaptic vesicles and the clamping of spontaneous neurotransmitter release rely on the integrity of the C-terminal half of the SNARE domain of syntaxin-1A. Elife 12, (2024).

71. Cao, Y. J., Wang, Q., Zheng, X. X., Cheng, Y. & Zhang, Y. Involvement of SNARE complex in the hippocampus and prefrontal cortex of offspring with depression induced by prenatal stress. J Affect Disord 235, 374–383 (2018).

72. Bradley, C. A. et al. A pivotal role of GSK-3 in synaptic plasticity. Front Mol Neurosci 5, (2012).

73. Jorge-Torres, O. C. et al. Inhibition of Gsk3b Reduces Nfkb1 Signaling and Rescues Synaptic Activity to Improve the Rett Syndrome Phenotype in Mecp2-Knockout Mice. Cell Rep 23, 1665–1677 (2018).

74. Duda, P., Hajka, D., Wójcicka, O., Rakus, D. & Gizak, A. GSK3β: A Master Player in Depressive Disorder Pathogenesis and Treatment Responsiveness. Cells 9, 727 (2020).

75. Nairn, A. C. & Shenolikar, S. The role of protein phosphatases in synaptic transmission, plasticity and neuronal development. Curr Opin Neurobiol 2, 296– 301 (1992).

76. Hayne, M. & DiAntonio, A. Protein phosphatase 2A restrains DLK signaling to promote proper Drosophila synaptic development and mammalian cortical neuron survival. Neurobiol Dis 163, 105586 (2022).

77. Iacob, E. et al. Dysregulation of leukocyte gene expression in women with medication-refractory depression versus healthy non-depressed controls. BMC Psychiatry 13, 273 (2013).

78. Szopa, A. et al. Purinergic transmission in depressive disorders. Pharmacol Ther 224, 107821 (2021).

79. Ren, J. & Bertrand, P. P. Purinergic receptors and synaptic transmission in enteric neurons. Purinergic Signal 4, 255–266 (2008).

80. Ortiz, R., Ulrich, H., Zarate, C. A. & Machado-Vieira, R. Purinergic system dysfunction in mood disorders: a key target for developing improved therapeutics. Prog Neuropsychopharmacol Biol Psychiatry 57, 117–131 (2015).

81. Jope, R. & Roh, M.-S. Glycogen Synthase Kinase-3 (GSK3) in Psychiatric Diseases and Therapeutic Interventions. Curr Drug Targets 7, 1421–1434 (2006).

82. Li, X., Liu, M., Cai, Z., Wang, G. & Li, X. Regulation of glycogen synthase kinase-3 during bipolar mania treatment. Bipolar Disord 12, 741–52 (2010).

83. Xu, L., Chen, Y., Shen, T., Lin, C. & Zhang, B. Genetic Analysis of PICK1 Gene in Alzheimer’s Disease: A Study for Finding a New Gene Target. Front Neurol 9, (2019).

84. Li, Y.-H., Zhang, N., Wang, Y.-N., Shen, Y. & Wang, Y. Multiple faces of protein interacting with C kinase 1 (PICK1): Structure, function, and diseases. Neurochem Int 98, 115–121 (2016).

85. Illes, P., Ulrich, H., Chen, J.-F. & Tang, Y. Purinergic receptors in cognitive disturbances. Neurobiol Dis 185, 106229 (2023).

86. Dev, K. K. & Henley, J. M. The schizophrenic faces of PICK1. Trends Pharmacol Sci 27, 574–579 (2006).

87. Fass, D. M. et al. Brain-specific deletion of GIT1 impairs cognition and alters phosphorylation of synaptic protein networks implicated in schizophrenia susceptibility. Mol Psychiatry 27, 3272–3285 (2022).

88. Kim, M. J. et al. Functional analysis of rare variants found in schizophrenia implicates a critical role for GIT1–PAK3 signaling in neuroplasticity. Mol Psychiatry 22, 417–429 (2017).

89. Alnafisah, R. et al. Altered purinergic receptor expression in the frontal cortex in schizophrenia. Schizophrenia 8, 96 (2022).

90. Salatino□Oliveira, A. et al. Association study of *GIT1* gene with attention□deficit hyperactivity disorder in Brazilian children and adolescents. Genes Brain Behav 11, 864–868 (2012).

91. Shim, S.-H. et al. Association between glycogen synthase kinase-3β gene polymorphisms and attention deficit hyperactivity disorder in Korean children: A preliminary study. Prog Neuropsychopharmacol Biol Psychiatry 39, 57–61 (2012).

92. Goehler, H. et al. A Protein Interaction Network Links GIT1, an Enhancer of Huntingtin Aggregation, to Huntington’s Disease. Mol Cell 15, 853–865 (2004).

93. Rippin, I. et al. Inhibition of GSK-3 ameliorates the pathogenesis of Huntington’s disease. Neurobiol Dis 154, 105336 (2021).

94. Varney, M. J. & Benovic, J. L. The Role of G Protein–Coupled Receptors and Receptor Kinases in Pancreatic β -Cell Function and Diabetes. Pharmacol Rev 76, 267–299 (2024).

95. Ding, H.-H., Ni, W.-J., Tang, L.-Q. & Wei, W. G protein-coupled receptors: potential therapeutic targets for diabetic nephropathy. Journal of Receptors and Signal Transduction 36, 411–421 (2016).

96. Murrell-Lagnado, R. D. Regulation of P2X Purinergic Receptor Signaling by Cholesterol. in 211–232 (2017). doi:10.1016/bs.ctm.2017.05.004.

97. Zhang, Q. et al. High Density Lipoprotein (HDL) Promotes Glucose Uptake in Adipocytes and Glycogen Synthesis in Muscle Cells. PLoS One 6, e23556 (2011).

98. Hirotani, S. et al. Inhibition of Glycogen Synthase Kinase 3β During Heart Failure Is Protective. Circ Res 101, 1164–1174 (2007).

99. Burnstock, G. Purinergic Signaling in the Cardiovascular System. Circ Res 120, 207–228 (2017).

100. Pang, J. et al. G-Protein–Coupled Receptor Kinase Interacting Protein-1 Is Required for Pulmonary Vascular Development. Circulation 119, 1524–1532 (2009).

101. Zhang, F., Armando, I., Jose, P. A., Zeng, C. & Yang, J. G protein-coupled receptor kinases in hypertension: physiology, pathogenesis, and therapeutic targets. Hypertension Research 47, 2317–2336 (2024).

102. Fang, K. C. Mesenchymal Regulation of Alveolar Repair in Pulmonary Fibrosis. Am J Respir Cell Mol Biol 23, 142–145 (2000).

103. Li, X.-W. et al. Role of eukaryotic translation initiation factor 3a in bleomycin-induced pulmonary fibrosis. Eur J Pharmacol 749, 89–97 (2015).

104. Yang, Y. et al. Screening for diagnostic targets in tuberculosis and study on its pathogenic mechanism based on mRNA sequencing technology and miRNA-mRNA-pathway regulatory network. Front Immunol 14, (2023).

105. Zhai, W., Wu, F., Zhang, Y., Fu, Y. & Liu, Z. The Immune Escape Mechanisms of Mycobacterium Tuberculosis. Int J Mol Sci 20, 340 (2019).

106. Krassnig, S. et al. A Profound Basic Characterization of eIFs in Gliomas: Identifying eIF3I and 4H as Potential Novel Target Candidates in Glioma Therapy. Cancers (Basel) 13, 1482 (2021).

107. Bertorello, J. et al. Translation reprogramming by eIF3 linked to glioblastoma resistance. NAR Cancer 2, (2020).

108. Braganhol, E., Wink, M. R., Lenz, G. & Battastini, A. M. O. Purinergic Signaling in Glioma Progression. in 87–108 (2020). doi:10.1007/978-3-030-30651-9_5.

109. Arcos-Montoya, D. et al. Progesterone Receptor Together with PKCα Expression as Prognostic Factors for Astrocytomas Malignancy. Onco Targets Ther Volume 14, 3757–3768 (2021).

110. Chorna, N. E. et al. P2Y _2_ receptors activate neuroprotective mechanisms in astrocytic cells. J Neurochem 91, 119–132 (2004).

111. Zhu, Y., et al. *PPP2R1B* Gene in Chronic Lymphocytic Leukemias and Mantle Cell Lymphomas. Leuk Lymphoma 41, 177–183 (2001).

112. Wu, X. et al. Targeting glycogen synthase kinase 3 for therapeutic benefit in lymphoma. Blood 134, 363–373 (2019).

113. Zhang, Q., Madden, N. E., Wong, A. S. T., Chow, B. K. C. & Lee, L. T. O. The Role of Endocrine G Protein-Coupled Receptors in Ovarian Cancer Progression. Front Endocrinol (Lausanne) 8, (2017).

114. Butler, M. S. et al. Small glutamine-rich tetratricopeptide repeat–containing protein alpha is present in human ovaries but may not be differentially expressed in relation to polycystic ovary syndrome. Fertil Steril 99, 2076–2083.e1 (2013).

115. Wu, R., Connolly, D. C., Ren, X., Fearon, E. R. & Cho, K. R. Somatic Mutations of the PPP2R1B Candidate Tumor Suppressor Gene at Chromosome 11q23 are Infrequent in Ovarian Carcinomas. Neoplasia 1, 311–314 (1999).

116. Zhang, Y. et al. eIF3a improve cisplatin sensitivity in ovarian cancer by regulating XPC and p27Kip1 translation. Oncotarget 6, 25441–25451 (2015).

117. Hu, L.-P. et al. Targeting Purinergic Receptor P2Y2 Prevents the Growth of Pancreatic Ductal Adenocarcinoma by Inhibiting Cancer Cell Glycolysis. Clinical Cancer Research 25, 1318–1330 (2019).

118. Elmadbouh, O. H. M., Pandol, S. J. & Edderkaoui, M. Glycogen Synthase Kinase 3β: A True Foe in Pancreatic Cancer. Int J Mol Sci 23, 14133 (2022).

119. Mai, W. et al. Glycogen synthase kinase 3β promotes osteosarcoma invasion and migration via regulating PTEN and phosphorylation of focal adhesion kinase. Biosci Rep 41, (2021).

120. Zhang, L. et al. Translational Regulation by eIFs and RNA Modifications in Cancer. Genes (Basel) 13, 2050 (2022).

121. Fang, Q. et al. A novel homozygous nonsense variant of STX2 underlies non-obstructive azoospermia in a consanguineous Chinese family. J Hum Genet (2024) doi:10.1038/s10038-024-01288-9.

122. Jin, J. et al. Multi-omics study identifies that PICK1 deficiency causes male infertility by inhibiting vesicle trafficking in Sertoli cells. Reproductive Biology and Endocrinology 21, 114 (2023).

123. Staudt, M. D., de Oliveira, C. V. R., Lehman, M. N., McKenna, K. E. & Coolen, L. M. Activation of NMDA Receptors in Lumbar Spinothalamic Cells is Required for Ejaculation. J Sex Med 8, 1015–1026 (2011).

124. Gur, S. et al. Management of Erectile Function by Penile Purinergic P2 Receptors in the Diabetic Rat. Journal of Urology 181, 2375–2382 (2009).

125. Goodarzi, M. O. et al. Small glutamine-rich tetratricopeptide repeat-containing protein alpha (SGTA), a candidate gene for polycystic ovary syndrome. Human Reproduction 23, 1214–1219 (2008).

126. Chang, W. et al. Adipocytes from women with polycystic ovary syndrome demonstrate altered phosphorylation and activity of glycogen synthase kinase 3. Fertil Steril 90, 2291–2297 (2008).

127. Singh, A. K. et al. Targeting the A3 adenosine receptor to prevent and reverse chemotherapy-induced neurotoxicities in mice. Acta Neuropathol Commun 10, 11 (2022).

128. Xu, T. et al. Uncovering the role of FOXA2 in the Development of Human Serotonin Neurons. Advanced Science 10, (2023).

129. Yang, Y. A Novel Progress of FXYD6 Structure and Functions. Peer Reviewed Journal of Forensic & Genetic Sciences 4, (2022).

130. Wurtman, J. Depression and weight gain: the serotonin connection. J Affect Disord 29, 183–192 (1993).

131. Matza, L. S., Phillips, G. A., Revicki, D. A., Murray, L. & Malley, K. G. Development and validation of a patient-report measure of fatigue associated with depression. J Affect Disord 134, 294–303 (2011).

132. Kursun, O., Yemisci, M., van den Maagdenberg, A. M. J. M. & Karatas, H. Migraine and neuroinflammation: the inflammasome perspective. J Headache Pain 22, 55 (2021).

133. Al Ghadeer, H. A. et al. Migraine Headache and the Risk of Depression. Cureus (2022) doi:10.7759/cureus.31081.

134. Ding, Q., Markesbery, W. R., Chen, Q., Li, F. & Keller, J. N. Ribosome Dysfunction Is an Early Event in Alzheimer’s Disease. The Journal of Neuroscience 25, 9171–9175 (2005).

135. Perini, G. et al. Cognitive impairment in depression: recent advances and novel treatments. Neuropsychiatr Dis Treat Volume 15, 1249–1258 (2019).

136. Tian, L., Ma, L., Kaarela, T. & Li, Z. Neuroimmune crosstalk in the central nervous system and its significance for neurological diseases. J Neuroinflammation 9, 594 (2012).

137. Marcolongo-Pereira, C. et al. Neurobiological mechanisms of mood disorders: Stress vulnerability and resilience. Front Behav Neurosci 16, (2022).

138. Poller, W. C. et al. Brain motor and fear circuits regulate leukocytes during acute stress. Nature 607, 578–584 (2022).

139. Pavlov, V. A. & Tracey, K. J. Neural circuitry and immunity. Immunol Res 63, 38–57 (2015).

140. Rahal, D. et al. Positive and negative emotion are associated with generalized transcriptional activation in immune cells. Psychoneuroendocrinology 153, 106103 (2023).

141. Goossens, J., Morrens, M. & Coppens, V. The Potential Use of Peripheral Blood Mononuclear Cells as Biomarkers for Treatment Response and Outcome Prediction in Psychiatry: A Systematic Review. Mol Diagn Ther 25, 283–299 (2021).

142. Zhang, M. & Huang, B. The multi-differentiation potential of peripheral blood mononuclear cells. Stem Cell Res Ther 3, 48 (2012).

143. Oshaghi, M., Kourosh-Arami, M. & Roozbehkia, M. Role of neurotransmitters in immune-mediated inflammatory disorders: a crosstalk between the nervous and immune systems. Neurological Sciences 44, 99–113 (2023).

144. Chen, C.-S., Barnoud, C. & Scheiermann, C. Peripheral neurotransmitters in the immune system. Curr Opin Physiol 19, 73–79 (2021).

145. Junger, W. G. Immune cell regulation by autocrine purinergic signalling. Nat Rev Immunol 11, 201–212 (2011).

146. Drevets, W. C., Wittenberg, G. M., Bullmore, E. T. & Manji, H. K. Immune targets for therapeutic development in depression: towards precision medicine. Nat Rev Drug Discov 21, 224–244 (2022).

147. Osimo, E. F. et al. Inflammatory markers in depression: A meta-analysis of mean differences and variability in 5,166 patients and 5,083 controls. Brain Behav Immun 87, 901–909 (2020).

148. Dias, H. D. et al. Integrative systems neuroimmunology reveals leukocyte-expressing PAX6 as a critical predictor of major depressive disorder. Preprint at 10.1101/2024.09.25.614771 (2024).

149. Xu, J., Wang, N., Luo, J. & Xia, J. Syntabulin regulates the trafficking of PICK1-containing vesicles in neurons. Sci Rep 6, 20924 (2016).

150. Manabe, R., Kovalenko, M., Webb, D. J. & Horwitz, A. R. GIT1 functions in a motile, multi-molecular signaling complex that regulates protrusive activity and cell migration. J Cell Sci 115, 1497–1510 (2002).

151. Krawczyk, M. et al. Expression of RAB4B, a protein governing endocytic recycling, is co-regulated with MHC class II genes. Nucleic Acids Res 35, 595– 605 (2006).

152. Hanley, J. G. PICK1: A multi-talented modulator of AMPA receptor trafficking. Pharmacol Ther 118, 152–160 (2008).

153. Perrin, L. et al. Rab4b controls an early endosome sorting event by interacting with the γ subunit of the clathrin adaptor complex 1. J Cell Sci (2013) doi:10.1242/jcs.130575.

154. Honer, W. G. et al. The synaptic pathology of cognitive life□. Dialogues Clin Neurosci 21, 271–279 (2019).

155. Chen, Y.-T., Lin, C.-H., Huang, C.-H., Liang, W.-M. & Lane, H.-Y. PICK1 Genetic Variation and Cognitive Function in Patients with Schizophrenia. Sci Rep 7, 1889 (2017).

156. Li, Y.-H., Zhang, N., Wang, Y.-N., Shen, Y. & Wang, Y. Multiple faces of protein interacting with C kinase 1 (PICK1): Structure, function, and diseases. Neurochem Int 98, 115–121 (2016).

157. Casad, M. E. et al. Cardiomyopathy Is Associated with Ribosomal Protein Gene Haplo-Insufficiency in *Drosophila melanogaster*. Genetics 189, 861–870 (2011).

158. Malinow, R. & Malenka, R. C. AMPA Receptor Trafficking and Synaptic Plasticity. Annu Rev Neurosci 25, 103–126 (2002).

159. Lim, B. K., Huang, K. W., Grueter, B. A., Rothwell, P. E. & Malenka, R. C. Anhedonia requires MC4R-mediated synaptic adaptations in nucleus accumbens. Nature 487, 183–189 (2012).

160. Pignatelli, M. et al. Cooperative synaptic and intrinsic plasticity in a disynaptic limbic circuit drive stress-induced anhedonia and passive coping in mice. Mol Psychiatry 26, 1860–1879 (2021).

161. Soda, T., Pasqua, T., De Sarro, G. & Moccia, F. Cognitive Impairment and Synaptic Dysfunction in Cardiovascular Disorders: The New Frontiers of the Heart–Brain Axis. Biomedicines 12, 2387 (2024).

162. Read, J. R., Sharpe, L., Modini, M. & Dear, B. F. Multimorbidity and depression: A systematic review and meta-analysis. J Affect Disord 221, 36–46 (2017).

163. Pizzol, D. et al. Relationship between severe mental illness and physical multimorbidity: a meta-analysis and call for action. BMJ Mental Health 26, e300870 (2023).

164. Berk, M. et al. Comorbidity between major depressive disorder and physical diseases: a comprehensive review of epidemiology, mechanisms and management. World Psychiatry 22, 366–387 (2023).

165. Hare, D. L., Toukhsati, S. R., Johansson, P. & Jaarsma, T. Depression and cardiovascular disease: a clinical review. Eur Heart J 35, 1365–1372 (2014).

166. Ghanei Gheshlagh, R., Parizad, N. & Sayehmiri, K. The Relationship Between Depression and Metabolic Syndrome: Systematic Review and Meta-Analysis Study. Iran Red Crescent Med J 18, (2016).

167. Bialek, K., Czarny, P., Strycharz, J. & Sliwinski, T. Major depressive disorders accompanying autoimmune diseases – Response to treatment. Prog Neuropsychopharmacol Biol Psychiatry 95, 109678 (2019).

168. Galts, C. P. C. et al. Depression in neurodegenerative diseases: Common mechanisms and current treatment options. Neurosci Biobehav Rev 102, 56–84 (2019).

169. Krawczyk, M. et al. Expression of RAB4B, a protein governing endocytic recycling, is co-regulated with MHC class II genes. Nucleic Acids Res 35, 595– 605 (2006).

170. Jacobson, K. A. et al. A _3_ Adenosine Receptors as Modulators of Inflammation: From Medicinal Chemistry to Therapy. Med Res Rev 38, 1031–1072 (2018).

171. Nishat, S., A Khan, L., M Ansari, Z. & F Basir, S. Adenosine A _3_ Receptor: A promising therapeutic target in cardiovascular disease. Curr Cardiol Rev 12, 18– 26 (2016).

172. Perrin, L. et al. Rab4b controls an early endosome sorting event by interacting with the γ subunit of the clathrin adaptor complex 1. J Cell Sci (2013) doi:10.1242/jcs.130575.

173. Incalza, M. A. et al. Oxidative stress and reactive oxygen species in endothelial dysfunction associated with cardiovascular and metabolic diseases. Vascul Pharmacol 100, 1–19 (2018).

174. Medina-Leyte, D. J. et al. Endothelial Dysfunction, Inflammation and Coronary Artery Disease: Potential Biomarkers and Promising Therapeutical Approaches. Int J Mol Sci 22, 3850 (2021).

175. Terashima, A. et al. An Essential Role for PICK1 in NMDA Receptor-Dependent Bidirectional Synaptic Plasticity. Neuron 57, 872–882 (2008).

176. Wu, A. & Zhang, J. Neuroinflammation, memory, and depression: new approaches to hippocampal neurogenesis. J Neuroinflammation 20, 283 (2023).

177. Pasquini, S. et al. Adenosine Receptors in Neuropsychiatric Disorders: Fine Regulators of Neurotransmission and Potential Therapeutic Targets. Int J Mol Sci 23, 1219 (2022).

178. Kang, J. et al. Ribosomal proteins and human diseases: molecular mechanisms and targeted therapy. Signal Transduct Target Ther 6, 323 (2021).

179. Jacovetti, C., Bayazit, M. B. & Regazzi, R. Emerging Classes of Small Non-Coding RNAs With Potential Implications in Diabetes and Associated Metabolic Disorders. Front Endocrinol (Lausanne) 12, (2021).

180. Wang, F. et al. Identification of differentially expressed genes of blood leukocytes for Schizophrenia. Front Genet 15, (2024).

181. Berk, M. et al. Comorbidity between major depressive disorder and physical diseases: a comprehensive review of epidemiology, mechanisms and management. World Psychiatry 22, 366–387 (2023).

