## Supplementary Figures for "Dysregulation of synaptic-related genes of neuroimmune networks within peripheral blood mononuclear cells in major depressive disorder"

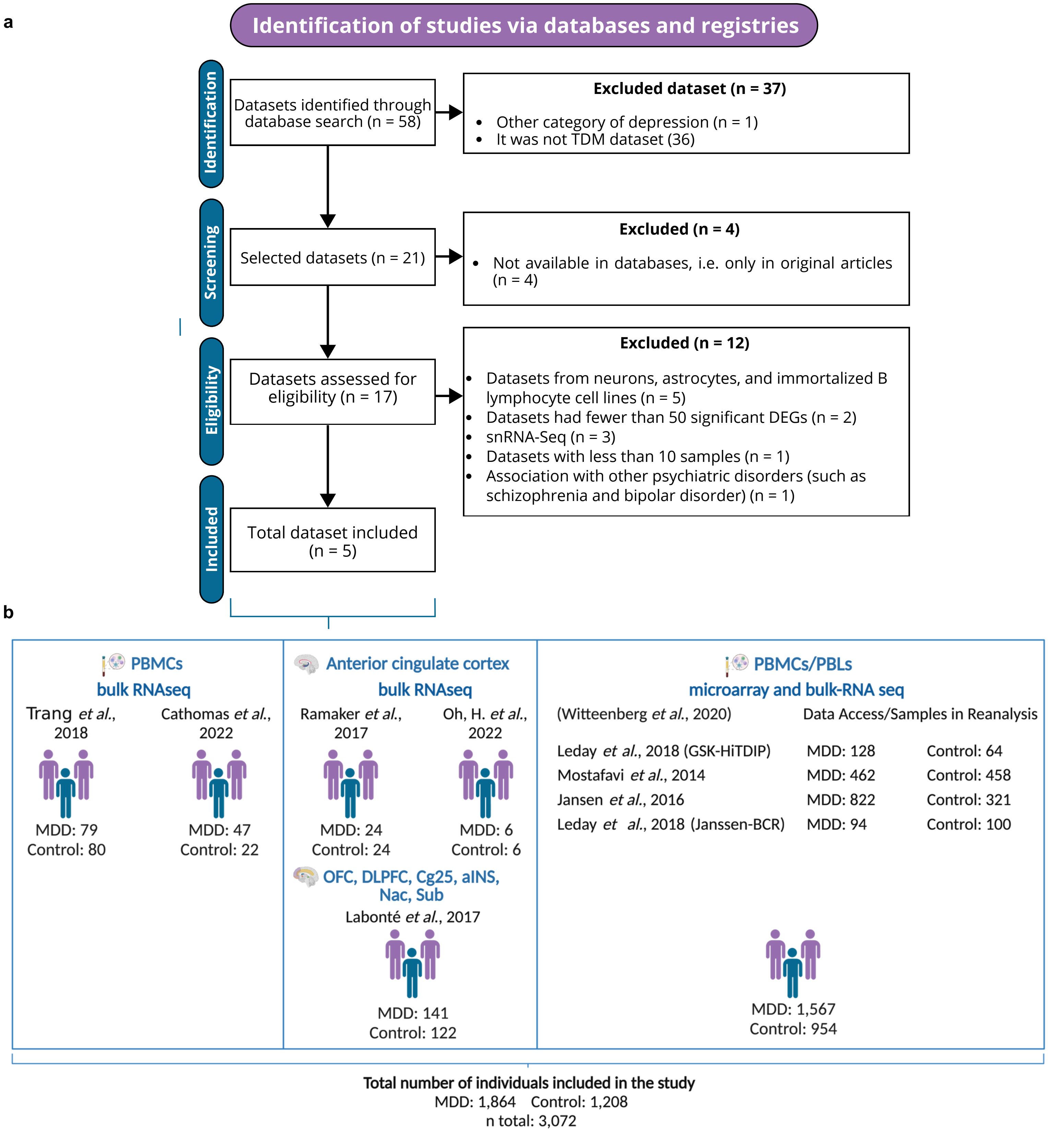


**Supplementary Figure 1. Flowchart of data curation.** (a) Workflow of the systematic literature search and dataset evaluation process. (b) Summary of datasets and samples included in the study, with details on sequencing methods, tissue origins, and sample sizes. A total of 3,072 samples was included, integrating data from Wittenberg et al. (2020) meta-analysis. This meta-analysis compiled findings from four large, independent case-control studies on whole blood samples, encompassing 1,567 cases of MDD and 954 healthy controls. Figure created with BioRender.

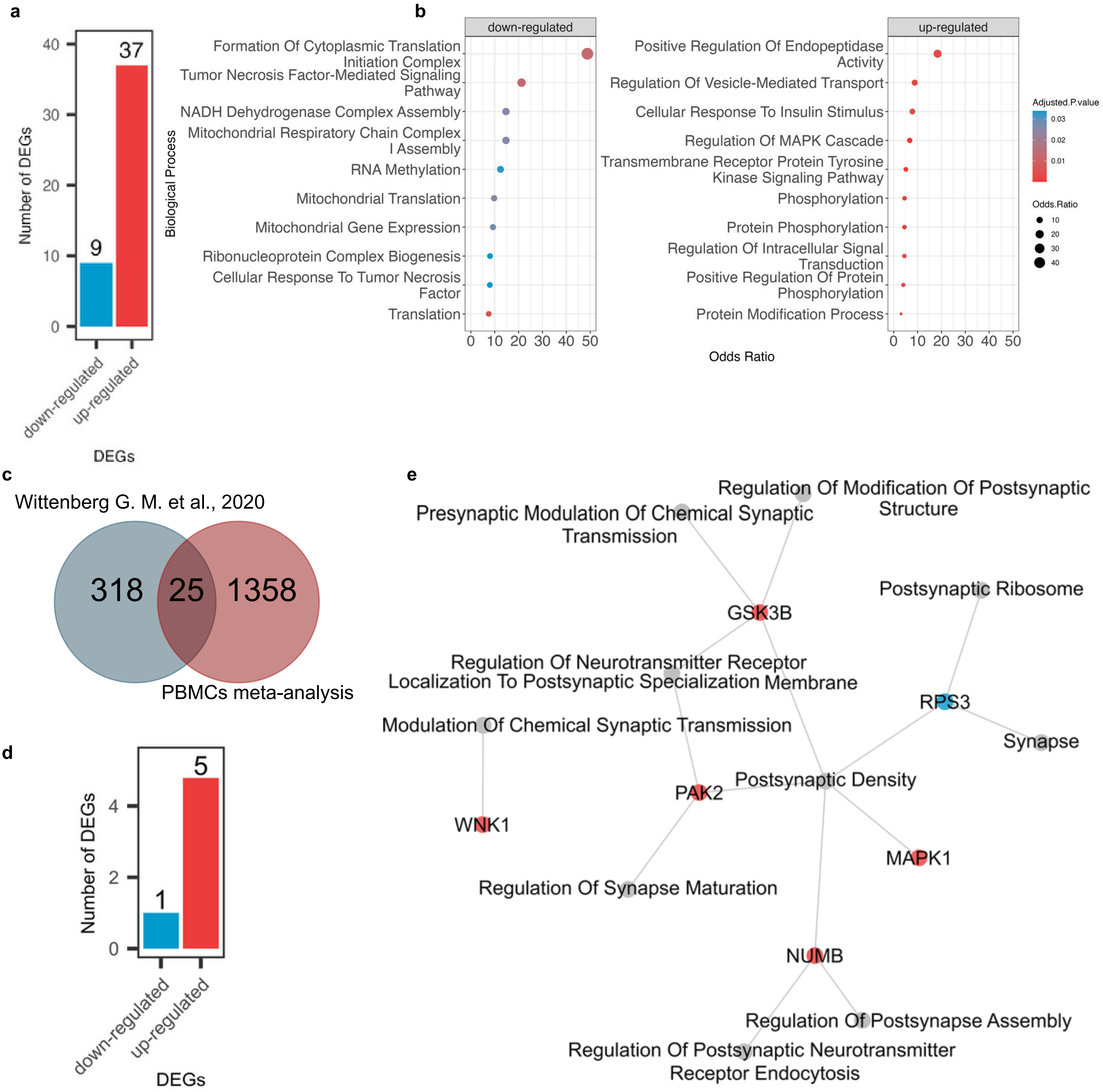


**Supplementary Figure 2. Comparison of the Wittenberg et al. (2020) study with the PBMC meta-analysis for synaptic ontology.** (a) Bar plot showing that out of 343 DEGs identified in the Wittenberg study, 46 were enriched for synaptic ontologies (comprising 9 down-regulated and 37 up-regulated metaDEGs). (b) Enrichment analysis of the 46 metaDEGs: down-regulated (left) and up-regulated (right). (c) Venn diagram illustrating the 25 common DEGs shared between the Wittenberg study and the PBMC meta-analysis conducted in our study. (d) Bar plot indicating that 6 of these 25 common metaDEGs were enriched for synaptic ontologies (1 down-regulated and 5 up-regulated). (e) Network representation of significant synaptic ontologies and the 6 DEGs.

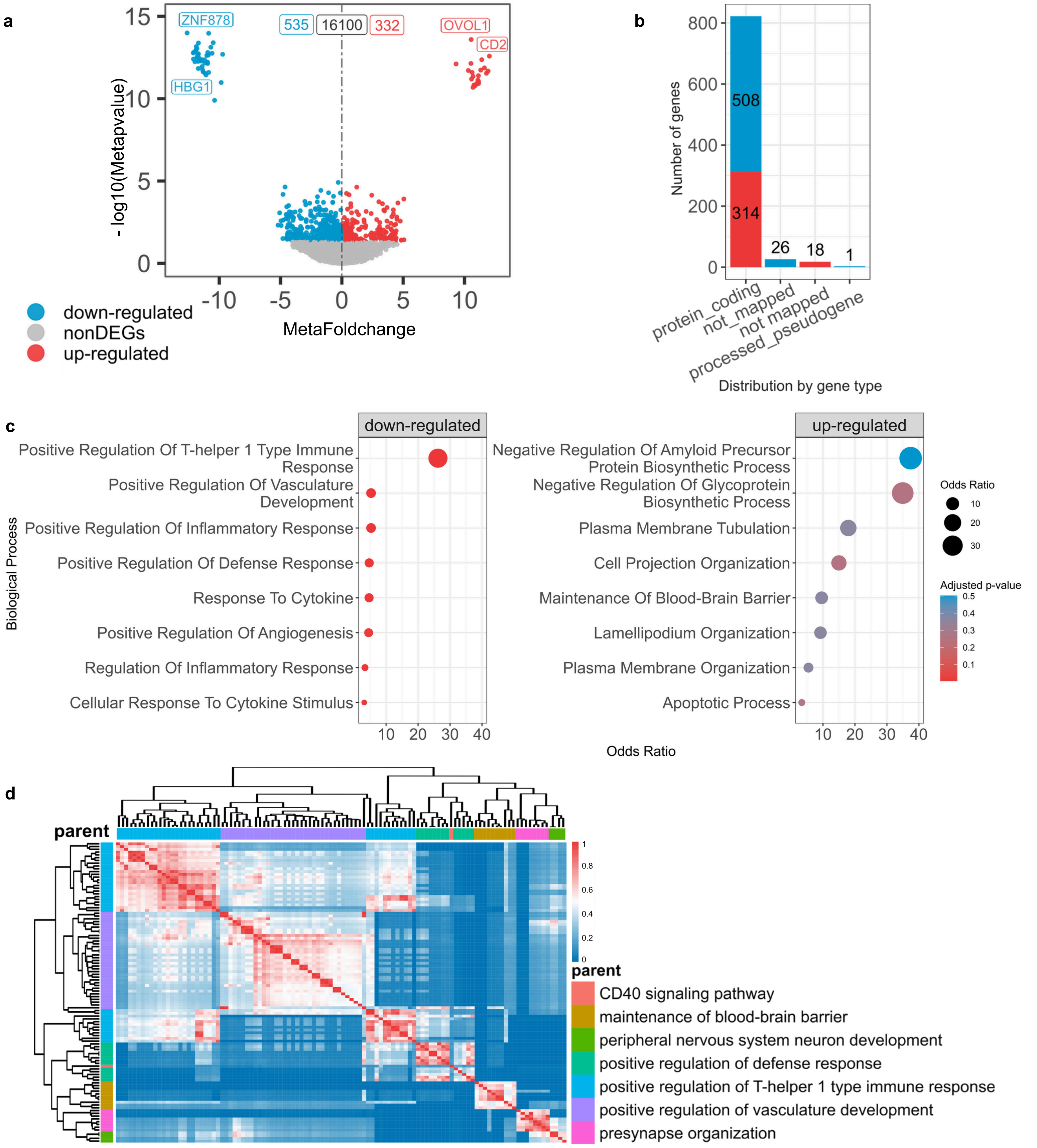


**Supplementary Figure 3. Meta-analysis and characterization of biological process interactions in the anterior cingulate cortex (ACC).** (a) Volcano plot displaying the distribution of metaDEGs. (b) Distribution of up-regulated (red) and down-regulated (blue) metaDEGs. (c) Enrichment analysis of metaDEGs, showing the results for down-regulated DEGs (left) and up-regulated DEGs (right). (d) Heatmap illustrating the similarity matrix of BPs derived from rrvgo analysis, highlighting the relationships among the enriched processes.
